# Truncated TrkB: The predominant TrkB Isoform in Nociceptors

**DOI:** 10.1101/2024.12.27.630513

**Authors:** Jaclyn Merlo, Chang Fang-Mei, Michael Tran, Jessie Alfaro, Tarek Ibrahim, Ping Wu, Shivani Ruparel

**Author notes:** **Corresponding author:** Shivani Ruparel, PhD, Center for Pain Therapeutics and Addiction Research, Department of Endodontics, University of Texas Health Science Center at San Antonio, 7703 Floyd Curl Drive, San Antonio, TX 78229.

## Abstract

Truncated TrkB (TrkBT1), traditionally considered a dominant-negative regulator of full-length TrkB (TrkBTK+), remains poorly understood in peripheral sensory neurons, particularly nociceptors. Furthermore, sensory neuronal TrkB expression and function has been traditionally associated with non-nociceptive neurons, particularly Aδ low-threshold mechanoreceptors. This study challenges prevailing assumptions by demonstrating that TrkBT1 is the predominant TrkB isoform expressed in sensory neurons and plays a functional role in modulating neuronal activity. We demonstrate that TrkBT1 is the predominant isoform expressed in nociceptors, identified by markers such as TRPV1, TRPA1, TRPM8 and 5HT3A, as well as non-nociceptors, while the full-length isoform (TrkBTK+) is restricted to non-nociceptive subpopulation. Functionally, we show that acute application of BDNF induces modest calcium influx in nociceptors and prolonged BDNF exposure significantly potentiates capsaicin-induced calcium influx, an effect blocked by the TrkB-specific antagonist ANA12. Additionally, BDNF also promotes the survival of both nociceptive and non-nociceptive neurons in culture, an effect dependent on TrkBT1 activity. Our data also reveal that ANA12 inhibits BDNF-mediated neuronal sensitization and survival in a concentration-dependent manner, implicating distinct TrkBT1 signaling pathways in these processes. Collectively, our findings redefine TrkBT1 as a functional modulator of nociceptor activity rather than a passive regulator of full-length TrkB. By uncovering its dual roles in nociceptor sensitization and survival, this study provides new insights into the molecular mechanisms of BDNF/TrkB signaling in pain. Future work evaluating the role of TrkBT1 in sensory biology could offer new perspectives on how this receptor contributes to neuronal function and plasticity during chronic pain conditions.

## Background

The brain-derived neurotrophic factor (BDNF) has been extensively studied in the context of pain and is known to contribute to various chronic pain conditions, including inflammatory[6; 15; 17; 29; 34], neuropathic[4; 26], orofacial[3], and cancer-related pain[1; 7; 12; 30]. As a member of the neurotrophin family, BDNF is expressed in both the peripheral and central nervous systems. A significant portion of research has focused on how BDNF influences pain through its actions at the synapses between peripheral sensory neurons and second-order neurons in the spinal cord. Under normal conditions, BDNF is constitutively expressed in the cell bodies of neurons in the dorsal root ganglia (DRG) and trigeminal ganglia (TG), and following injury, its expression increases and anterogradely transported to the central terminals to enhance post-synaptic excitability, contributing to chronic pain states[19; 26; 33].

Beyond neurons, BDNF is also reported to be produced by non-neuronal cells such as glial cells and immune cells in response to inflammatory signals and injury to contribute to synaptic plasticity and central sensitization[19; 26; 33]. Recently, we and others reported that BDNF can also be produced at the peripheral site of injury. For instance, our research demonstrated that BDNF is released by oral cancer cells in tongue tumors of patients, as well as in preclinical models, and modulates peripheral cancer-induced pain transmission and transduction[7; 12]. Similarly, *Gowler* et al. found that BDNF is present in the synovial fluid of osteoarthritis (OA) patients and mouse models, where it contributes to peripheral nociceptive behaviors[11]. Taken together, BDNF signaling axis plays a substantial role in promoting pain transmission, transduction and processing in a broad range of chronic pain states and is thus considered highly valuable in obtaining insights into potential targets for pain therapy.

It is widely accepted and demonstrated that BDNF produces its effects by specifically binding to its receptor TrkB leading to its activation and downstream signaling to modulate pain. TrkB is a tyrosine kinase receptor and exists in two spliced variants: 1) the full-length or long isoform (TrkBTK+) that signals via its kinase domain, and 2) the truncated or short isoform (TrkBT1) devoid of the kinase domain but harboring a unique 11 amino-acid sequence in the intracellular domain[2; 4; 9; 28]. TrkBTK+ is a widely studied and well characterized isoform, whereas TrkBT1 is highly understudied due to the initial assumptions that it acts as a dominant negative to the long form. However, it was later shown that BDNF can potentially signal through TrkBT1 independently of the long form, implying its role beyond being a dominant negative[2; 4; 9; 28]. Despite extensive research on BDNF signaling in pain, the role of TrkBT1 in nociceptive mechanisms is unclear. Till date, there are only three publications reporting the role of TrkBT1 in pain and these reports demonstrate its role in the central mechanism of pain in the spinal cord, particularly in glial cells[18; 23; 32]. Therefore, a peripheral role of this isoform is not investigated till date, despite evidence suggesting that TrkBT1 may play a significant role in BDNF-mediated peripheral pain. For example, the study by *Gowler* et al demonstrated that TrkBT1 is the predominant isoform expressed in the synovial fluid during OA and intraarticular sequestration of TrkB reversed OA-mediated pain[11]. Moreover, our prior study revealed that oral cancer-released BDNF contributed to cancer-induced pain via sensory neuronal activation of TrkB[12]. Importantly, we showed that TrkBT1 is the predominant isoform, expressed in tongue-innervating trigeminal neurons, compared to the full-length TrkB, across both peptidergic as well as non-peptidergic neuronal subgroups[12]. Building upon these findings, the current study aimed to further characterize TrkB isoforms in trigeminal neurons and evaluate the functional relevance of this isoform in BDNF-mediated activation and survival of trigeminal sensory neurons. Our data revealed that TrkBT1 is the predominant isoform in the trigeminal nociceptor and non-nocicpetor subpopulation and mediates BDNF-induced activation, sensitization and survival of nociceptors.

## Materials and Methods

### Animals

Six-to eight-week-old adult C57BL6 male and female mice (Jackson Labs, Bar Harbor, ME, USA) were used for most experiments. We also bred Rosa26^LSL-tDTTomato+^(Jackson Labs) with CGRP^cre+/-ER^ to obtain CGRP-ER-TDT male mice expressing TDT reporter in CGRP+ neurons for a few experiments. CGRP^cre+/-ER^ mice were provided by Dr. Armen Akopian at UTHSCSA, who obtained these mice from Dr. Pao-Tien Chuang (UC San Francisco, San Francisco, CA). CGRP-ER-TDT mice were induced by injecting 100mg/kg tamoxifen (Sigma-Aldrich, St. Louis, MO) intraperitoneally, three times, every other day. All animal handling and procedures were performed according to approved UTHSCSA IACUC protocols and conformed to the guidelines of International Association for the Study of Pain (IASP). The study is reported in accordance with the ARRIVE guidelines. Purchased animals were housed in the UTHSCSA laboratory of Animal Resources (LAR) for at least 4 days prior to start of experiments.

### Drugs

Recombinant BDNF was purchased from PeproTech (Cranbury, NJ, USA, catalog Number 450-02) and diluted to 100ng/ml stock in 5% Trehalose in sterile water The TrkB receptor antagonist, ANA-12 was purchased from Sigma Aldrich (St. Louis, MO, USA) and diluted to 12mg/ml in 100% DMSO. This stock was further diluted to desired concentration in media or PBS. Capsaicin powder was purchased from Sigma Aldrich and diluted at 100mM stock in ethanol to further dilute in PBS.

### Western Blot

Trigeminal ganglia were collected from naïve male and female mice and processed in 200 µL of RIPA buffer supplemented with a cOmplete Mini protease inhibitor cocktail (Millipore Sigma, St. Louis, MO). The tissue was homogenized using a Qiagen TissueLyser (Bio-Rad, Hercules, CA, USA) set to 50 oscillations per second for 1 minute. This step was followed by a 15-minute incubation on ice and repeated once. The samples were centrifuged at 14,000 × g for 10 minutes at 4 °C, and the resulting supernatant was collected as cell lysates for western blot analysis. Protein concentrations in the lysates were determined using the Pierce™ BCA Protein Assay Kit (ThermoFisher Scientific, Carlsbad, CA), following the manufacturer’s instructions. Equal protein amounts from each sample were resolved on 4–20% Mini-PROTEAN® TGX Stain-Free™ Protein Gels (12-well, 20 µL capacity) using SDS-PAGE, according to the Bio-Rad protocol. Proteins were then transferred to LF PVDF membranes using the Trans-Blot® Turbo™ Transfer System (Bio-Rad). After transfer, membranes were stained with Ponceau S to confirm protein transfer, washed three times with deionized water, and blocked for 1 hour at room temperature with a solution of 50 mL Tris-buffered saline with Tween 20 (TBST) and 2.5 g skim milk powder (Sigma-Aldrich, St. Louis, MO). Membranes were then incubated overnight at 4 °C with primary antibodies diluted in the blocking solution: anti-TRKB (1:500, Cell Signaling Technology, Cat# 4607S) and anti-GAPDH (1:5000, Cell Signaling Technology, Cat# 14C10). Following primary antibody incubation, membranes were washed three times for 10 minutes each with 0.1 M TBST and incubated for 1 hour at room temperature with secondary antibodies (1:5000, Amersham ECL HRP-Linked IgG, Cat# NA934-100UL, Cytiva) diluted in blocking solution. Membranes were washed again three times for 10 minutes each with 0.1 M TBST. Proteins were visualized using the WesternBright™ ECL detection kit (Advansta, San Jose, CA), and chemiluminescence was captured using a Bio-Rad ChemiDoc imaging system. Densitometry analysis of the protein bands was performed using ImageJ software.

### Retrograde labeling

Isoflurane inhalation was used to briefly anesthetize mice to retrolabel tongue innervating trigeminal neurons. Animals were injected with 5ul of 0.5% WGA-488 (ThermoFisher Scientific, Carlsbad, CA) as described previously[21]. To ensure thorough labeling of the lingual neurons, WGA was injected twice per side of the tongue—once at the surface and once into deeper layers of the tissue.

### Single Cell RTPCR

Two days after the WGA injections, trigeminal ganglion (TG) tissues were dissected and dissociated in a sterile petri dish. Fluorescently labeled neurons were manually isolated using a Narishige micromanipulator and FERTY Syringe Plus Microinjector under an inverted EVOS FL digital fluorescence microscope (AMG, Bothell, WA). Neurons were categorized into three size groups: small (<15 µm), medium (<25 µm), and large (>25 µm). Each single cell was placed in 4 µL of 2X reaction mix from the CellsDirect™ One-Step qRT-PCR Kit (ThermoFisher Scientific, Carlsbad, CA). Target genes were pre-amplified using Fluidigm PreAmp master mix and TaqMan probes (Applied Biosystems). A 200 pg sample of Universal Mouse RNA (BioChain, Newark, CA) served as the positive control, while a no-template control (NTC) was used as the negative control. Additionally, a PreAmp NTC was included to ensure the quality of primer amplification. Gene expression analysis (qRT-PCR) was conducted on 48.48 IFCs (integrated fluidic circuits) using the BioMark™ HD system (Fluidigm, Inc.) following the manufacturer’s instructions. Ct values for each gene in each cell were normalized to the internal control gene UBB. Relative gene expression levels were calculated using the 2^(-delta Ct) method.

### Primary Culture of TG neurons

Trigeminal ganglia (TGs) were dissected and prepared for neuronal cultures as described previously[14; 24]. Tissues were washed in HBSS and then dissociated using 1 mg/mL collagenase and dispase (Roche, Indianapolis, IN, USA) for 45 minutes at 37°C. The dissociated cells were plated on poly-D-lysine/laminin-coated glass coverslips (BD Biosciences, San Jose, CA, USA) and cultured in DMEM media supplemented with glutamine, penicillin/streptomycin, and 2% fetal bovine serum for the vehicle group. For the experimental group, the same media was used, supplemented with 100 ng/mL recombinant BDNF (rBDNF) and/or the selected concentration of ANA12. ANA12 was added to the cultures two hours before rBDNF. Cultures were grown overnight for calcium imaging experiments and for 48 hours for immunohistochemical staining.

### Calcium Imaging

Calcium accumulation in cultured sensory neurons was assessed using fluorescence imaging, as described previously[14; 24]. TG cultures were incubated with the calcium-sensitive dye Fura-2 AM (2 µM; Molecular Probes, Carlsbad, CA, USA) in Hank’s modified buffer. Imaging was conducted with a Nikon TE 2000U microscope equipped with a ×40/1.35 NA Fluor objective. Neurons were monitored for calcium influx following drug applications, including 100 ng/mL BDNF, 10nM capsaicin, or 20 mM potassium chloride (KCL). The net change in calcium influx was determined by subtracting the average baseline intracellular calcium concentration [Ca2+]i from the peak [Ca2+]i observed after exposure to capsaicin.

### Immunohistochemistry

Immunohistochemistry was performed on both mouse TG tissues and TG neuronal cultures. TG tissues were collected from anesthetized mice following intracardiac perfusion with 4% paraformaldehyde (PFA). The TGs were dissected, post-fixed in 4% PFA for 20 minutes, washed in 0.1M phosphate buffer (PB), and cryosectioned at a thickness of 20 µm using Neg-50 (Richard Allan, Kalamazoo, MI, USA). The tissue sections were then subjected to immunostaining as previously described[12-14]. Briefly, sections were washed in 0.1M phosphate-buffered saline (PBS) and incubated for 90 minutes at room temperature in a blocking solution consisting of 4% normal goat serum (Sigma, St. Louis, MO), 2% bovine gamma-globulin (Sigma-Aldrich, St. Louis, MO), and 0.3% Triton X-100 (Fisher Scientific) in 0.1M PBS. The sections were then incubated with primary antibodies diluted in the blocking solution for 16 hours at room temperature. The primary antibodies used were: 1:150 anti-TrkB (Cell Signaling), 1:200 NeuN (Neuromics), and 1:300 TRPV1 (Neuromics). Following incubation with the primary antibodies, sections were washed and incubated with secondary antibodies in blocking solution for 90 minutes at room temperature. Secondary antibodies were purchased from Molecular Probes, Eugene, OR, USA. After staining, the tissue sections were washed in 0.1M PBS and deionized water, air-dried in the dark, and mounted with Vectashield Antifade Mounting Medium with DAPI (Vectorlabs, Burlingame, CA, USA). For the staining of neuronal cultures, TG neurons were grown on coverslips, post-fixed with 4% PFA for 10 minutes, followed by a 15-minute treatment with 100% ethanol, and then washed with PBS containing 1.2 mM calcium chloride to prevent cell loss during staining. The cells were then blocked and stained using the protocol described above. Images were captured using a Keyence microscope (BZ-X800E) equipped with a 20× objective and identical laser gain settings across all experimental groups. The laser gain settings were calibrated to ensure no-positive staining in the no-primary control. Images were acquired using fixed acquisition parameters for all groups. For quantitative analysis, six images were taken from each coverslip, and the total number of neuronal cells in these images was counted across all experimental groups.

### Statistics

Statistical analysis was conducted using GraphPad Prism 8.0 (GraphPad, La Jolla, CA). Sample sizes were planned with G*Power, based on effect sizes and standard deviations derived from preliminary data, to achieve 80% statistical power at a two-sided alpha level of 0.05. The results are shown as the mean ± SEM. Group comparisons were made using either an unpaired Student’s t-test or one-way/two-way ANOVA, with Bonferroni correction applied as needed. A two-sided alpha level of 0.05 was used to determine statistical significance.

## Results

### Truncated TrkB is the predominant isoform expressed in trigeminal neurons including nociceptors

In our previous work[12], we established that truncated TrkB is the dominant TrkB isoform in trigeminal neurons. In this study, we examine its expression across both sexes and further investigated its presence within the nociceptor subset of trigeminal sensory neurons. As illustrated in **Fig 1A and B**, truncated TrkB remained the predominant isoform in trigeminal ganglia (TG) tissue from both male and female subjects, with similar expression levels between the sexes. To assess TrkB expression in nociceptors, we evaluated its colocalization within TRPV1-expressing neurons. **Fig 2C** shows that TrkB indeed colocalizes with several TRPV1-positive neurons, suggesting its expression in nociceptors. However, these findings provide qualitative, rather than quantitative, evidence of TrkB in nociceptors, and the antibody used does not differentiate between TrkB isoforms. To confirm TrkBT1 expression specifically in the nociceptor population, we analyzed single-cell RT-PCR data from retrolabeled tongue-innervating TG neurons. Approximately 35% of sensory neurons expressed TrkBT1, compared to ∼11% that expressed the full-length TrkBTK+ isoform (**Fig 1D and E**). We further quantified TrkBT1 presence in various nociceptor and non-nociceptor subpopulations using specific markers. As shown in **Fig 1F**, TrkBT1 was found in approximately 22.5% of TRPV1+ neurons, 33.5% of TRPA1+ neurons, 31% of TRPM8+ neurons, and 47% of 5HT3A+ nociceptors. Conversely, TrkBTK+ was present in roughly 2% of TRPV1+ neurons, 5% of TRPA1+ neurons, 4% of TRPM8+ neurons, and was absent in 5HT3A+ neurons. In non-nociceptor subtypes, both TrkBT1 and TrkBTK+ were equally expressed in ∼55% of Parv+ neurons whereas 50% of TrkC+ neurons expressed TrkBT1, while ∼13% of TrkC+ neurons expressed TrkBTK+ **(Fig 1G)**. Collectively, these results confirm that TrkBT1 is the predominant isoform in trigeminal sensory neurons, with its expression found in both nociceptor and non-nociceptor subpopulations of trigeminal sensory neurons. In contrast TrkBTK+ is expressed in a small proportion of sensory neurons, largely restricted to non-nociceptor populations.

**Fig 1.**
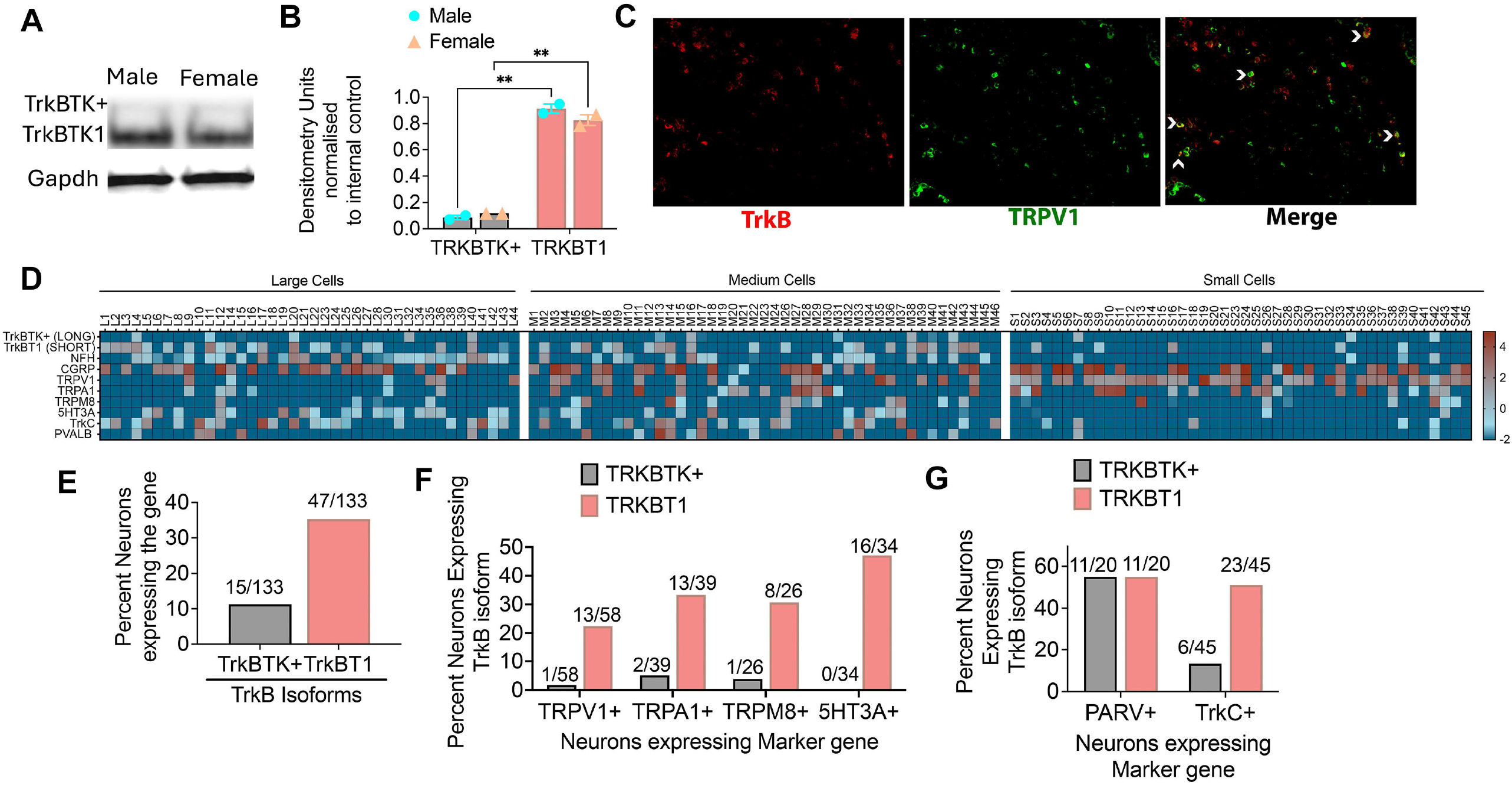
Expression of TrkB Isoforms in Trigeminal Sensory Neurons. **A**. TrkB isoform protein expression in naïve male and female mice. TrkBTK+ is shown at 125kDa and TrkBT1 is shown at 90kDa. **B**. Densitometry analyses of Western blot data. n=3 per group. Data represented as mean±SEM and analyzed by two way ANOVA with Sidak post hoc test at p<0.05. **C**. Cryosections of TG tissues from naïve mice were stained with TrkB and TRPV1. Arrows indicate colocalization. Images taken at 20x magnification. **D**. Mice were injected with 0.5% WGA in the tongue and 2 days later, WGA-positive small, medium, and large cells were subjected to single-cell RTPCR for the expression of TrkB long (TrkBTK+), short TrkB (TrkBT1), CGRP, NFH,TRPV1, TRPA1, TRPM8, 5HT3A, TrkC and Parvalbumin. Data represented as heatmap of relative gene expression normalized to internal control. **E**. Percentage of neurons expressing TrkB isoforms. Percentage of neurons expressing TrkB isoforms **F**. in nociceptors and **G**. in non-nociceptors.

### BDNF-induces calcium influx in nociceptors

Given the specific expression of TrkBT1 in nociceptors, we investigated its functional role by examining whether BDNF induces calcium influx in trigeminal neurons. BDNF application elicited a modest yet consistent calcium influx (**Fig. 2A, 2B**) in 43% of all recorded trigeminal sensory neurons (**Fig. 2C**). To further assess whether BDNF directly modulates nociceptor activity, we analyzed calcium responses in capsaicin-positive (CAP+) neurons. As shown in **Fig. 2D** and **2E**, BDNF triggered calcium influx in CAP+ neurons. Additionally, capsaicin responses were observed in approximately 35% of cultured neurons (**Fig. 2F**), and among these, ∼23% also exhibited BDNF-induced calcium influx (**Fig. 2G**). Together, these findings demonstrate the functional relevance of BDNF signaling in modulating nociceptor activity.

**Fig 2.**
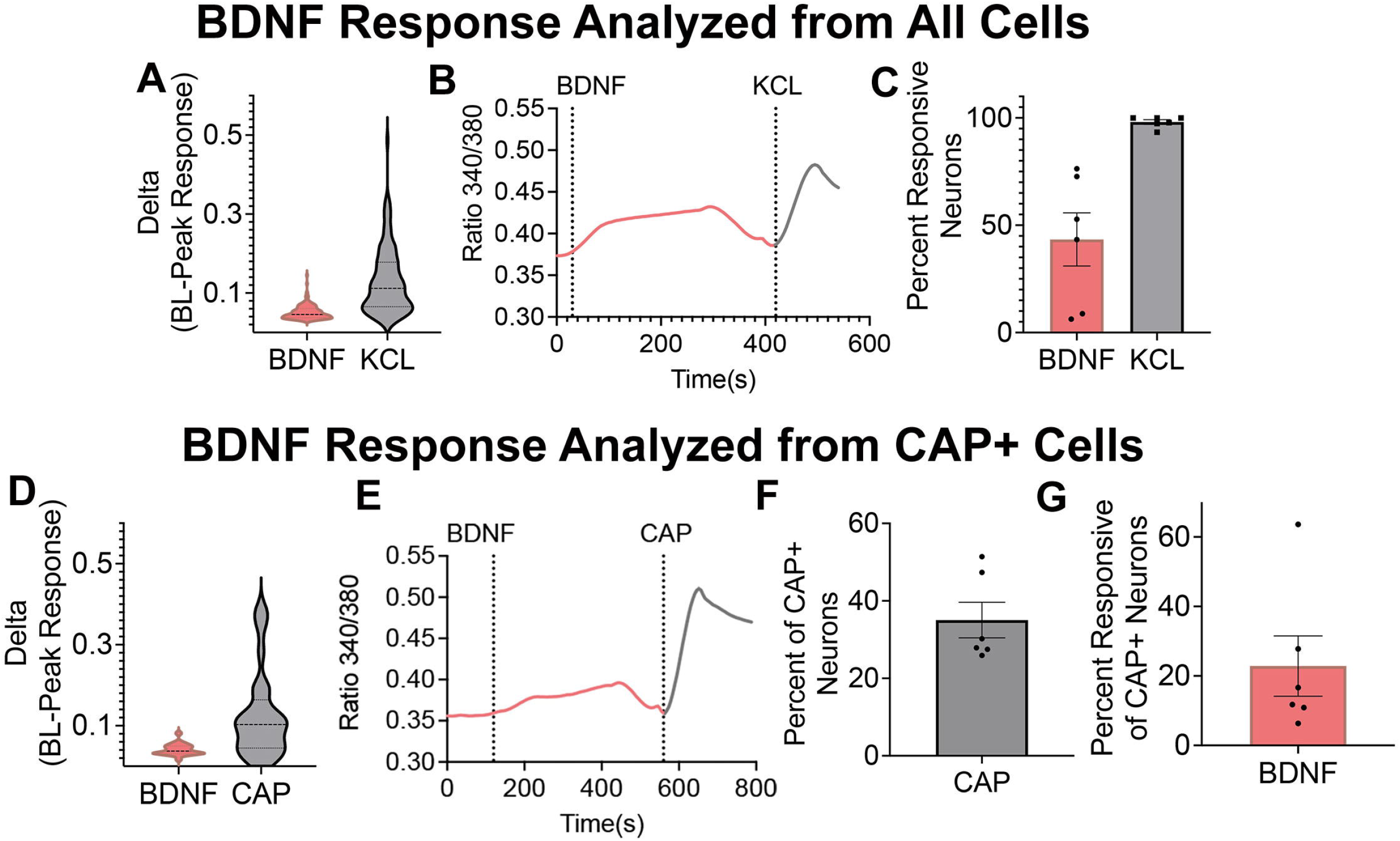
Effect of BDNF on calcium accumulation in nociceptors. **A-C**. Mouse TG neurons were cultured overnight and 100ng/ml BDNF was applied for 5 mins, followed by washout period and application of 20mM potassium chloride (KCL). **A**. BDNF response plotted as delta 340/380 ratio from baseline. **B**. Representative traces of BDNF-induced [Ca^2+^]i is shown. **C**. Percent cell responsive to BDNF and KCL. **D-G:** Mouse TG neurons were cultured overnight and 100ng/ml BDNF was applied for 5 mins, followed by washout period and application of 10nM capsaicin (CAP). **D**. BDNF response plotted as delta 340/380 ratio from baseline. **E**. Representative traces of BDNF-induced [Ca^2+^]_i_ in CAP+ neurons is shown. **F**. Percent cell responsive to CAP. **G**. Percent of BDNF responsive cells among all CAP+ cells. Data represented as mean±SEM.

### BDNF potentiates TRPV1 responses via Truncated TrkB

In addition to investigating the direct activation of nociceptors by BDNF, we also assessed whether BDNF sensitizes the trigeminal nociceptor population via TrkB. Acute application of BDNF to sensory neurons did not affect CAP-evoked calcium accumulation **(Fig 3A and B)** nor did it alter the number of CAP-responsive neurons **(Fig 3C)**. However, prolonged incubation of neuronal cultures with BDNF resulted in a dose-dependent potentiation of CAP-evoked calcium accumulation **(Fig 3D, E)** without changing the proportion of CAP-responsive neurons **(Fig 3F)**. Additionally, co-treatment with ANA12 prevented the BDNF-induced sensitization of sensory neurons to CAP **(Fig 4)**. ANA12 binds to TrkB at two distinct sites with different IC50 values. The high-affinity binding site inhibits BDNF activity at an IC50 of 45.6nM while the low-affinity site does so at an IC50 of 41.1uM. To determine which of these sites binding sites regulates BDNF-mediated nociceptor activity, we tested the effects of ANA12 at 100nM and 10µM. As shown in **Fig 4A and B**, 100nM ANA12 significantly, yet partially inhibited the BDNF-potentiated CAP response, while a complete inhibition was observed with the 10 µM dose. ANA12 treatment had no effect on proportion of CAP-responsive neurons **(Fig 4C)**. These findings suggest that BDNF may regulate nociceptor activity by binding to both sites on TrkBT1.

**Fig 3.**
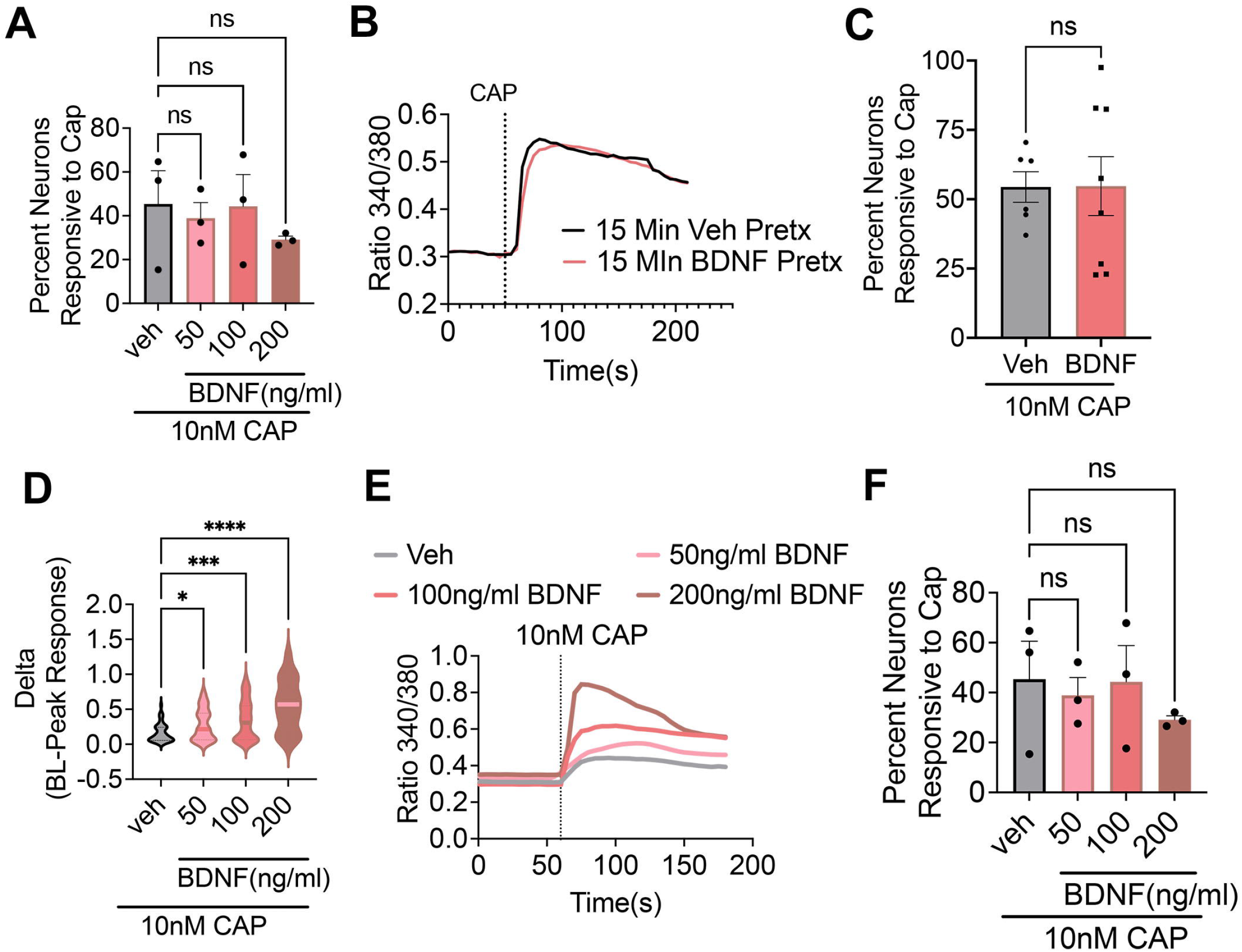
Effect of BDNF on TRPV1 sensitization in sensory neurons. **A-C**. Mouse TG neurons were cultured overnight and at the time of recording, cells were treated with veh or 100ng/ml BDNF for 15 mins prior to application of 10nM Capsaicin for 2 mins. **A**. Cap response plotted as delta 340/380 ratio from baseline. **B**. Representative traces of CAP-induced [Ca^2+^]_i_ is shown. **C**. Percent cell responsive to CAP. **D-F**. Mouse TG neurons were cultured overnight with veh or recombinant BDNF at 50, 100 or 200ng/ml. Cells were tested for [Ca^2+^]_i_ upon 10nM capsaicin (CAP) application for 2mins. **D**. Cap response plotted as delta 340/380 ratio from baseline. **E**. Representative traces of CAP-induced [Ca^2+^]_i_ is shown. **F**. Percent cell responsive to CAP. Data represented as mean±SEM and analyzed with 1-way ANOVA with the Sidak post hoc test at P<0.05.

**Fig 4.**
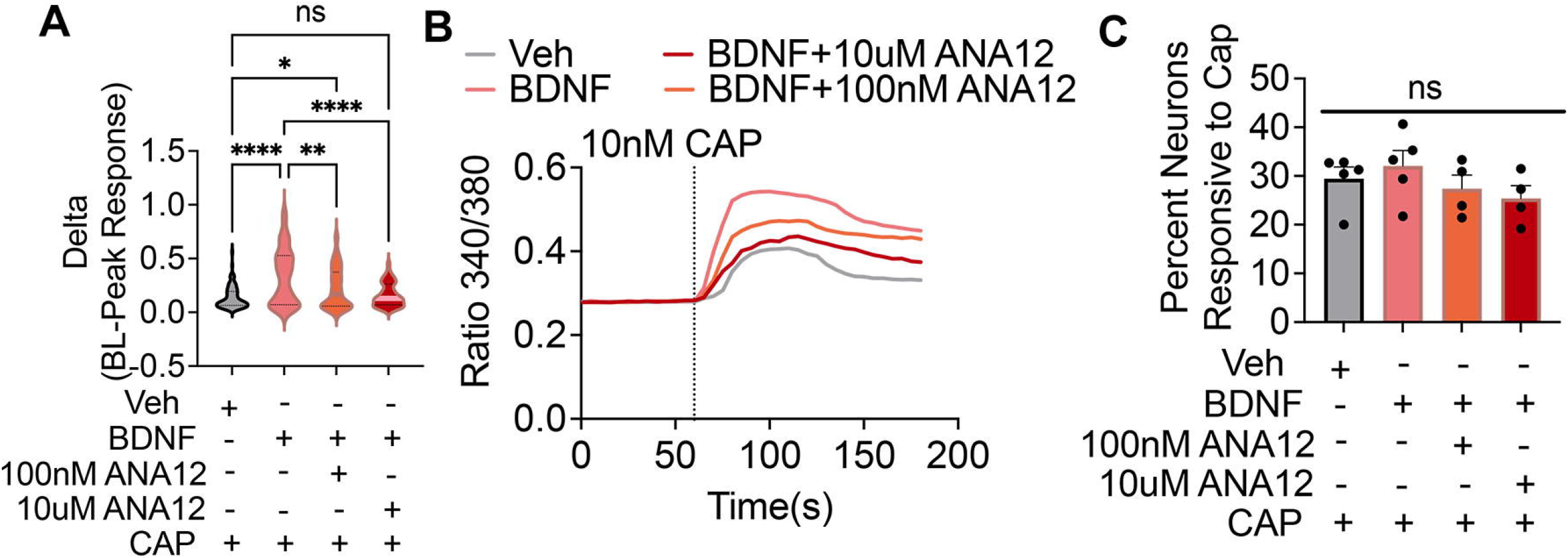
Effect of TrkB inhibition on BDNF-mediated sensitization of TRPV1+neurons. Mouse TG neurons were cultured with either veh/veh, Veh/100ng/ml BDNF or 100nM ANA12/100ng/ml BDNF or 10uM ANA12/100ng/ml BDNF or 24 hours. Following this, cells were tested for [Ca^2+^]_i_ following 10nM CAP for 2 mins. **A**. Cap response plotted as delta 340/380 ratio from baseline. **B**. Representative traces of CAP-induced [Ca^2+^]_i_ is shown. **C**. Percent neurons responsive to CAP. Data represented as mean±SEM and analyzed with 1-way ANOVA with the Sidak post hoc test at P<0.05.

### Truncated TrkB contributes to BDNF mediated increase in survival of nociceptor and non-nociceptor population of trigeminal sensory neurons

Given BDNF’s trophic abilities, we next tested whether BDNF/TrkBT1 axis contributes to survival of trigeminal neurons in culture. To differentiate between nociceptor and non-nociceptor subpopulation, we cultured neurons from CGRP-TDT mice and stained with a pan-neuronal marker NeuN. Total number of neurons per slide as well as number of CGRP+ and CGRP-neurons were quantified in cultures treated with vehicle, BDNF and BDNF with three doses of ANA12. Our data revealed that BDNF treated cultures had significantly greater number of NeuN+ neurons compared to vehicle treated group (**Fig 5A and B**) confirming prior finding that BDNF increases survival of sensory neurons[22]. Additional analyses showed that both CGRP+ and CGRP-neurons were increased with BDNF treatment (**Fig 5 A, C, D**). These data indicate that BDNF may improve survival of nociceptor as well as non-nociceptor subtypes of sensory neurons. Furthermore, treatment of ANA12 dose dependently inhibited BDNF-mediated increase in survival of all NeuN+ neurons (Fig 5A, B) as well as CGRP+ **(Fig 5A and C)** and CGRP-neurons (**Fig 5A and D**). Given that TrkBT1 is the predominant isoform in all trigeminal neurons and the only isoform expressed in nociceptors, these data indicate that BDNF increase in sensory neuronal survival is TrkBT1 dependent. Importantly, doses of ANA12 required to inhibit neuronal survival were found to be higher than that needed for neuronal sensitization.

**Fig 5.**
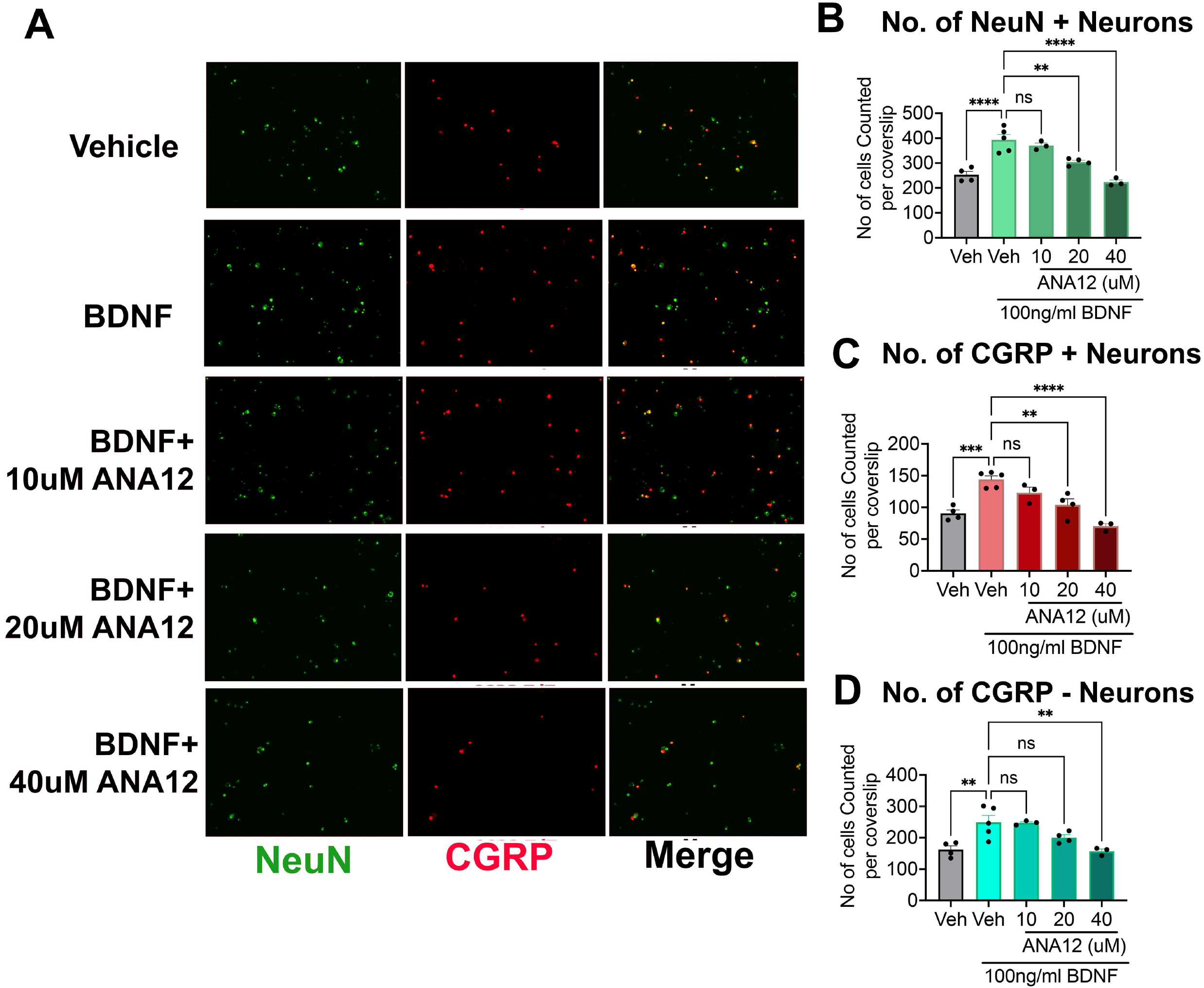
Effect of BDNF on survival of trigeminal sensory neurons in culture. CGRP-ER-TDT mice were induced with Mouse TG neurons harvested from CGRP-ER-TDT mice were cultured with either veh/veh, Veh/100ng/ml BDNF , 10uM ANA12/100ng/ml BDNF, 20uM ANA12/100ng/ml BDNF or 40uM ANA12/100ng/ml BDNF for 24 hours. Cells were stained with pan-neuronal marker NeuN and imaged using Keyence microscope with 10x magnification and counted for NeuN+, CGRP+ and CGRP-neurons per image. **A**. Representative image of culture shown for each group. **B**. Total Number of NeuN+ cells counted for each group per coverslip. **C**. Total Number of CGRP+ neurons counted for each group per coverslip. **D**. Total Number of CGRP-neurons counted for each group per coverslip. Data represented as mean±SEM and analyzed with 1-way ANOVA with the Sidak post hoc test at P<0.05.

## Discussion

Our study investigates the expression and functional relevance of TrkBT1 in trigeminal nociceptors. This work was built upon our previous work as well as the study by *Gowler* et al demonstrating that peripheral BDNF signaling via TrkB plays a pivotal role in chronic pain conditions such as oral cancer -induced pain[7; 12] and osteoarthritis[11] respectively. Additionally, our study revealed that the truncated isoform of TrkB is the predominant isoform expressed across a wider proportion of trigeminal sensory neurons compared to the full-length receptor[12]. While the role of TrkBT1 in astrocytes and glial cells has been explored for central pain mechanisms[18; 23; 32], its relevance in peripheral nociception remained unclear. Given the critical role of nociceptors in pain modulation, we aimed to characterize the expression and function of TrkBT1 in this neuronal subpopulation.

Our results demonstrated that nociceptors, identified by markers such as TRPV1, TRPA1, TRPM8, and 5HT3A, predominantly express TrkBT1. Conversely, the non-nociceptor subpopulation, characterized by parvalbumin and TrkC, expresses both isoforms of TrkB. These data challenges the longstanding view that TrkB is confined to non-nociceptor populations, particularly the Aδ-LTMR subtype[16; 31]. Our findings suggest that while TrkBTK+ is likely to be restricted to Aδ non-nociceptors, TrkBT1 constitutes a significant portion of the total TrkB-expressing population and may play a pivotal role in modulating neuronal activity during pain.

To confirm the functional relevance of TrkBT1, we examined whether BDNF can induce calcium influx in sensory neurons. Acute BDNF application elicited modest calcium influx in sensory neurons, including capsaicin-responsive neurons, suggesting that BDNF modulates nociceptor activity. However, in our prior study we reported that exogenous BDNF injections in naïve mice tongue tissue failed to induce pain behaviors[12], indicating that this modest calcium increase is insufficient to evoke pain under naïve conditions. Instead, prior neuronal priming (e.g., during injury) to enhance TrkBT1 expression or activity, maybe required to enable BDNF to produce pain. To this end, our data also showed that acute application of BDNF was unable to affect capsaicin responses from neurons cultured from naïve animals. Interestingly though, long-term exposure of cells to BDNF (i.e condition likely to be mimicked during injury or pathological states), dose dependently produced potentiation of capsaicin-induced calcium influx, that was inhibited by TrkB-specific antagonist ANA12. This finding indicates that perhaps the modest but sustained calcium influx by the interaction of BDNF and TrkBT1 may drive downstream changes in neuronal plasticity and ion channel modulation, such as sensitization of TRPV1.

Interestingly, ANA12 demonstrated differential effects depending on concentration: nanomolar doses partially inhibited BDNF sensitization, whereas micromolar doses completely blocked it. Given that ANA12 binds TrkB at both high- and low-affinity sites at IC50 values of 45.6nM and 41.1uM respectively[5], these findings imply that its effects on TrkBT1 occur primarily at the low-affinity binding site in sensory neurons.

Beyond nociceptor sensitization, our study also explored the neurotrophic effects of BDNF. Using transgenic mice expressing TdTomato in CGRP+ neurons, to distinguish between nociceptor and non-nociceptor population, we showed that BDNF promotes survival of both CGRP+ and CGRP-neurons in culture suggesting that BDNF mediates its neurotrophic effects on all sensory neuronal subtypes. ANA12 dose-dependently inhibited this effect, indicating that TrkBT1, rather than TrkBTK+, plays a predominant role in BDNF-mediated sensory neuron survival.

Interestingly, our survival assay revealed that BDNF-TrkBT1 interactions may alter the cytoskeleton to promote cell attachment, as survival was measured by the number of neurons attached to glass coverslips, aligning with previous studies implicating TrkBT1 in regulating the actin cytoskeleton and RhoA signaling[20]. However, since BDNF also sensitizes nociceptors via TrkBT1, this suggests that multiple downstream mechanisms are involved. This is further supported by our observation that the ANA12 concentrations required to inhibit neuronal sensitization differ from those needed to inhibit cell survival, indicating mechanistically distinct pathways. Specifically, doses needed to inhibit cell survival were higher than those needed to inhibit neuronal sensitization. This is an important finding and supports the idea that if TrkBT1 were to be targeted for pain relief, pharmacological inhibition of the receptor could modulate pain pathways without detrimental effects on neuronal health.

Taken together, our findings highlight, for the first time, the active role of TrkBT1 in nociceptor activation, sensitization, and survival. These results challenge the traditional view of TrkBT1 as a mere dominant-negative regulator of full-length TrkB. Instead, they suggest that TrkBT1 plays a critical role in modulating neuronal activity and survival, opening new avenues for understanding its molecular underpinnings in chronic pain and for developing targeted therapies.

## Limitations and Future Directions

One limitation of our study is the use of the pan-TrkB antagonist ANA12, which inhibits both TrkBT1 and full-length TrkB. Although TrkBT1’s predominant expression in nociceptors minimizes confounding effects, future studies should employ more targeted approaches, such as TrkBT1-specific siRNA/shRNA or knockout models. However, designing siRNA/shRNA is challenging due to the sequence overlap between TrkBT1 and full-length TrkB. Leveraging previously developed TrkBT1 global knockout or floxed mouse lines could overcome this hurdle. Additionally, small molecules that selectively inhibit TrkBT1 without affecting full-length TrkB would be valuable tools for both research and therapeutic applications.

While BDNF is known to promote axonogenesis and neurite outgrowth[8; 10; 25; 27], our study did not examine TrkBT1 trafficking to peripheral nerve terminals or its role in BDNF-mediated neurite outgrowth. Future work should address these aspects, particularly under chronic pain conditions, to elucidate the full spectrum of TrkBT1 functions. Finally, exploring how injury-induced changes in TrkBT1 expression or activity contribute to heightened neuronal plasticity and overall pain pathophysiology could deepen our understanding of its role in peripheral pain mechanisms and TrkBT1 biology. Furthermore, our prior study demonstrating that inhibition of neuronal TrkB reverses BDNF-mediated oral cancer pain behaviors[12], warrants an investigation on whether TrkBT1 can be used as a potential target for chronic pain treatment.

## Acknowledgements

Study was supported by funds provided by R01DE029187(SR), R01DE027223 (SR) R56DE034087 and T32 DE014318-23 COSTAR Program. All this support was provided by the National Institute of Dental and Craniofacial Research. Single-cell gene expression analysis was performed by Bioanalytic and Single-cell core (BASiC) at UT Health at San Antonio. The BASiC core is supported by the Cancer Prevention Research Institute of Texas (RP150600) and the Office of Vice President of Research of UT Health at San Antonio.

## Author Contributions

JM performed majority of calcium imaging and immunohistochemistry experiments as well as analyzed the data and made figures. Additionally, she also assisted in preparation of other experiments, lead the overall project and coordinated with others to enable timely completion of all experiments. CFM conducted a portion of calcium imaging experiments. MT conducted western blot experiments. TI conducted a portion of IHC staining. JA conducted all breeding and genotyping for the study. PW conducted single-cell RTPCR experiments. S.R. acquired all images. S.R. conceptualized and designed all experiments. S.R and JM wrote the manuscript. All authors edited the manuscript.

## Conflict of Interest

Authors declare no Conflict of Interest

## Notes

### Competing Interest Statement

The authors have declared no competing interest.

